# Mycolic Acid like lipids act as substrates for *Mycobacterium tuberculosis melH*

**DOI:** 10.1101/2025.04.06.646916

**Authors:** Shreyoshi Chakraborti, Jyothi S Sistla

## Abstract

*Mycobacterium tuberculosis* (*Mtb*), the pathogenic bacterium that causes tuberculosis, has developed its own ways of evading defense mechanisms to counteract the lethal effects of reactive oxygen species (ROS) generated within the host macrophages during infection. The *melH* gene present in *Mtb* and *Mycobacterium marinum* (*Mm*) plays an important role to reduce ROS generated during infection. The *melH* gene encodes for an epoxide hydrolase. Bioinformatics data suggests that encoded enzyme utilizes lipid substrates for its function. Initially, we used a lipid fractionation approach coupled with liquid chromatography mass spectrometry (LC-MS) and treatment with active MelH enzyme to identify potential substrates for MelH. We found classes of mycolic acids, predominantly epoxy mycolic acids accumulate in the *melH* mutant and upon treatment with MelH are reduced in the lipid fraction. These results provide insight into how MelH encoded in the *mel2* operon contributes to *Mtb* virulence and persistence and present further evidence for potential mechanisms of action if MelH is targeted for antitubercular drug discovery.

## Introduction

Mycobacterium tuberculosis (*Mtb*) causes tuberculosis (TB), a major global health threat and the second leading cause of death from a single infectious agent after COVID-19 (1). The pathogen’s metabolic plasticity is the key to its success and a major reason for its resistance to a plethora of antibiotics (2). *Mtb* is able to survive in harsh cellular conditions and under nutrient starvation (3). One of the major stresses that the mycobacterium withstands is reactive oxygen stress (ROS). Several genes have been identified that contribute to the mycobacterium’s ability to resist oxidative stress (4). Cirillo et al have identified the *mel2* locus as important for conferring resistance to ROS (5). This *mel2* locus consists of 6 genes (*melF-melK*) and a *melH* (Rv1938) deletion mutant has the highest levels of ROS under peroxide stress compared to other mutants in both *Mtb* and *Mm* (5).

Recent evidence suggests MelH is an epoxide hydrolase (EphB) which converts epoxides to dihydrodiols (6). However the physiological substrate of this protein is still unknown. Bioinformatic studies suggest *melH* is similar to *lux* operon genes and is likely to utilize fatty acid substrates (7). *Mtb* H37Rv strain has six potential epoxide hydrolases (EHs), proposed to have a role in detoxification of xenobiotics and biotransformation of endogenous epoxides (8). Knocking out *melH* in *Mtb* or *Mm* results in increased ROS susceptibility, decreased ATP levels, accumulation of aldehydes, and redox imbalance (9).

In this study, we combined a liquid chromatography-mass spectrometry (LC-MS) method with lipidomic analyses to identify potential lipid classes that may act as physiological substrates of MelH. Our results suggest that MelH treatment leads to a decreased abundance of oxygenated mycolic acid lipids as established through detailed lipidomic analysis. Thus, epoxidized mycolic acids may serve as substrates for MelH. Overall, the structural elucidation of these lipid compounds opens avenues for understanding *Mtb’s* mechanisms of action against oxidative stress.

### Experimental Materials and Methods

#### General Methods and Materials

All chemicals and reagents were of analytical reagent grade and were procured from commercial sources. MelH was prepared as previously described (10). C17 Ceramide (C18:1/17:0) was obtained from Avanti Polar Lipids. Affinity Purification Buffer A: 50mM Tris HCl, 300mM NaCl pH 7.5 Buffer B: 50mM Tris HCl 50 mM NaCl pH 7.5. Ion Exchange Chromatography Buffer: 50 mM Tris HCl, 50-300 mM NaCl pH 7.5. MelH Activity Buffer: 50 mM Tris HCl pH 7.5

#### Bacterial strains and culture conditions

*Mm* strains were grown at 30 °C in Middlebrook 7H9 medium (Difco) supplemented with 0.2% glycerol or 0.1 mM cholesterol, 0.5% bovine serum albumin, 0.08% NaCl, and 0.05% (vol/vol) tyloxapol (Sigma-Aldrich). The medium was filter sterilized. Alternatively, *ΔmelH Mm*, ymm1 strains were grown in the presence of 50 µg/mL hygromycin B. To determine the effect of the carbon source on the MelH competition assay, Mm were cultured in Middlebrook 7H9 medium supplemented with 10% oleate-albumin-dextrose-NaCl (OADC), 0.05% tyloxapol, and one of the following carbon sources: glycerol, sodium propionate, or cholesterol to a final concentration of 50 µM.

#### Lipid Extraction

*Mm* WT (*ymm1*) and *ΔmelH Mm* strains were grown in propionate growth media and a modified Bligh and Dyer method (11) was used to prepare separate crude lipid extracts from the cell pellets and from the culture filtrates. Briefly, wet cells (200 mg) were pelleted from exponential growth cultures (OD-600 = 0.4), and the culture filtrate was separated from the cells. The cell pellets and culture filtrates were incubated separately for 20 min with 5 mL of hexane and then the hexane layer was removed. CHCl_3_-methanol-water (2:2:1; v/v/v, 6 mL) was added to the cell pellet and the culture filtrate separately after removal of the hexane layer and the mixture incubated for 2 days at 4 °C with occasional stirring to allow the phases to separate. For complete phase separation, the samples were centrifuged at 3000g for 10 min. The monophasic CHCl_3_ lipid extract was transferred to a 20 mL glass tube, and the extract dried under vacuum, and stored at -20 °C. Prior to LC-MS analysis, the dried lipid extract was dissolved in CHCl_3_ to approximately 4 µg/µL and transferred to an autosampler vial. The samples were normalized to the total amount of lipid for all carbon source comparisons.

#### Competition Assay

Dried lipid samples were redissolved in 5% dimethyl sulfoxide (DMSO) to achieve a final 1 mg/ml concentration. MelH activity was measured using Epoxy Fluor 7 (cyano(6-methoxy-2-napthalenyl) methyl[(2,3)-3-phenyloxiranyl]methyl ester, carbonic acid) as substrate (Cayman Chemical, USA) in 20 mM Tris-HCl (pH 7.5) containing 0.1 M NaCl at 30 °C in the dark. For the competition assay, the reaction mixture contained 150 nM MelH, 100 µM Epoxy Fluor 7 substrate, and 1 mg/ml lipid extracts, keeping the DMSO concentration at 1% (v/v) in the total reaction volume. For all reactions, the mixture was incubated for 15 min, and the formation of the fluorescent product was monitored at λ _Ex/Em_ = 360/485 nm. A negative control was included.

#### Enzyme Assay of Fractionated Lipid Classes

Lipid samples were divided into three equal 50 μl aliquots. All subsequent treatments were performed in technical triplicate. To fractionate the glyco, phospho, and hydrophobic lipids such as trehalodimycolates (TDM) and trehalomonomycolate (TMM)), the initial KO propionate culture filtrate total lipid extract (20 mg) was separated using a silica column (99 x 7) mm with approximately 0.6 g of silica gel (12). Before loading, the silica column was washed with CHCl_3_. Then the lipid sample was loaded onto the column in a volume of 5 mL. A step gradient was used to elute the lipids. Step 1. Diethyl ether:petroleum ether (98:2, v/v, 10 mL). Step 2. CHCl_3_:MeOH (95:5, v/v, 10 mL). Step 3. CHCl_3_:MeOH (85:15, v/v, 10 mL). Step 4. CHCl_3_:MeOH (75:25, v/v, 10 mL). Steps 2-4 provided complex lipids. 2 mL fractions were collected and each fraction was divided into two equal aliquots by volume, and the solvent evaporated. One fraction was dissolved in 50 μl 5% DMSO/95% 20 mM Tris HCl pH 7.5. The second fraction was dissolved in 50 μl 5% DMSO/95% Tris HCl pH 7.5 containing MelH to provide a final enzyme concentration of 5 µM, and the mixture was incubated at 37 °C for 3 h. The enzyme reaction and the control solution were both quenched with 50 μl CHCl_3_. The CHCl_3_ layers were separated and dried in vacuo and stored at -20 °C until analyzed. 1 µg of C17 ceramide (18:1/17:0) was added to each fraction as an internal standard. The samples were redissolved in CHCl_3_ immediately before injection for LC-MS analysis.

#### Lipidomic Sample Preparation and Data Collection

Samples were analyzed using an Agilent 1290 Infinity II system equipped with a series of columns (i) a Zorbax Eclipse Plus C18 column (1.8 μm, 2.1 x 50 mm), (ii) an ACQUITY UPLC BEH C18 column (1.7 μm, 2.1 x 100 mm, Waters), and (iii) a HiPlex H column (4.6 x 250 mm, Agilent). The column temperature was set to 45 °C with a binary solvent system and a flow rate of 200 μL/min. Reversed-phase separation was achieved using a gradient from solvent A (35% acetonitrile, 35% water, 30% methanol, and 10 mM ammonium formate) to solvent B (95% isopropanol, 5% water, 10 mM ammonium formate) as follows: 100% A held for 0–2 min to 100% B for 50–55 min, 100% A for 2 min, and re-equilibration for 3 min (total run time: 60 min/sample). The flow rate was maintained at 200 μL/min for the duration of the run; the injection volume was 10 μL; the column was held at 45°C; and samples were held at 4 °C. Samples were dissolved in 30 μl CHCl_3_ to a final concentration of 15 ng/μl. The column eluate was infused into a Bruker Impact II QTOF system with an electrospray ion source. Data were collected in positive ion mode, with the following settings: capillary voltage of 4,500 V; endplate offset of 500 V; nebulizer gas pressure of 1.8 bar; dry gas flow rate of 8.0 L/min; dry temperature of 220 °C; MS spectra acquisition rate of 8 Hz; and m/z range of 150–2600 Da.

#### Analysis of Lipidomic Data

The data were analyzed using Bruker Data Analysis software. Total ion chromatograms and extracted ion chromatograms were generated with a tolerance of <5 ppm. Ion chromatograms were aligned using XCMS (13). Molecular formulas and fragmentation trees were generated through Sirius 4 (14). The total ion chromatograms were normalized to the C17 ceramide internal standard. The intensities of total ion peaks for the MelH-treated sample were reduced between retention times (RT) of 3-12 min. In contrast, there was little variation in the total ion peak intensities between pre and post treatment in the 20-25 min RT region. Therefore, as a negative control, the fold change (pre/post treatment intensity) was calculated for m/z values in the 20-25 min RT region to set a lower cutoff for significance. This cutoff was 2-fold. Then, the ions in the 3-12 min RT region were analyzed up to two decimal points of mass accuracy. For each mass, the intensity and fold change (pre/post treatment intensity) were determined. Only masses that showed a fold change greater than 2 were analyzed further.

Molecular formulas were annotated through Bruker Metaboscape software by deconvoluting ion adducts, and individual metabolite monoisotopic masses were calculated and compared with the dataset using the *Mtb* LipidDB, Mycomap and HMDB libraries (15,16,17). The average peak intensities and standard deviations from three biological replicates were calculated. Additionally, mycolic acid lipid standard data were analyzed using both Data Analysis and Metaboscape software by Bruker to compare and annotate the untargeted lipidomic dataset.

## Results

As a first step to identify under what conditions the maximal amount of potential MelH substrates are accumulated, lipid extracts from cultures of *Mm* grown in three different carbon sources were analyzed. We performed a fluorometric synthetic substrate enzymatic assay in the presence of lipid extracts from cells and culture filtrates grown in glycerol, propionate and cholesterol to determine which fraction had the highest level of competition of fluorescent substrate turnover, and therefore most potential MelH substrate present. The lipid extract from *ΔmelH Mm* culture filtrate grown on propionate showed the highest level of competition with MelH activity. Most of the cellular lipid extracts did not compete with MelH turnover except for *ΔmelH* lipid extracts grown in cholesterol as a carbon source. Although lipids from cholesterol cellular lipids and culture filtrate both showed high competition of MelH fluorescent substrate turnover, *ΔmelH* propionate culture filtrate lipid extracts showed the highest amount of competition and therefore we selected propionate culture filtrate lipid extracts from *ΔmelH Mm* for our future experiments. (Fig 1) (18).

**Fig 1:**
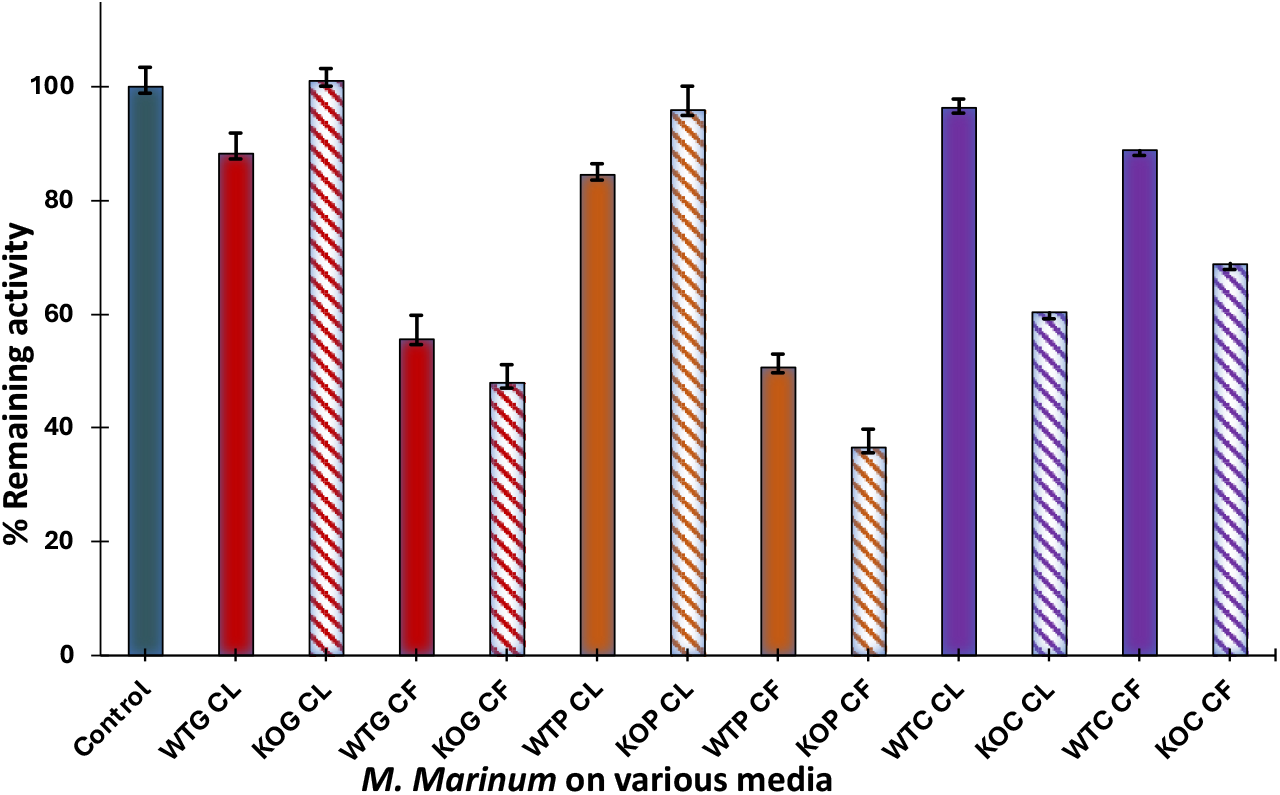
Competitive inhibition assay in presence of Epoxy Fluor7 and lipid extracts. 50 μM of artificial substrate was incubated with 150 ng of active MelH and 1μg/μl lipid extracts at pH 7.5, 37 °C for 15 mins. The control is the reaction with no lipid and the relative inhibition is compared with the negative control. CL indicates cellular lysate lipids and CF indicates Culture Filtrate lipids. G, P, C indicates glycerol, propionate and cholesterol carbon sources respectively. WT and KO represents WT *ymm1 Mm* strains and *ΔmelH Mm* strains respectively. The solid colors represent WT *Mm* strain and the lipids are extracted from different carbon source growth conditions whereas the striped represents *ΔmelH Mm* strain and the lipids extracted from different carbon source growth conditions.

The lipid extracts of the *ΔmelH Mm* propionate culture filtrate were fractionated using a silica column with a step gradient of chloroform and methanol. Each fraction of lipid was then treated with active MelH enzyme and the LC-MS total ion chromatograms of the fractions before and after treatment were compared. A significant decrease in abundance of ions was observed for peaks with retention times between 3 to 5 mins and 10 to 12 mins for the CHCl_3_:MeOH (85:15 v/v) fraction. However, no corresponding increase of ions potentially related to the enzymatic product were observed. The early retention time under reversed phase conditions suggested that the substrate is a hydrophilic lipid. Ceramide17 synthetic lipid (18:1/17:0) was used as an internal standard to normalize spectra. Verification of normalization was performed by confirming there were no changes in ion intensity in the 20-25 minute retention time region pre and post treatment. Additional analysis of lipid extracts treated with heat denatured MelH protein was also performed as a negative control (Fig 2). Spectra of all other fractions are available in Fig S4.

**Fig 2:**
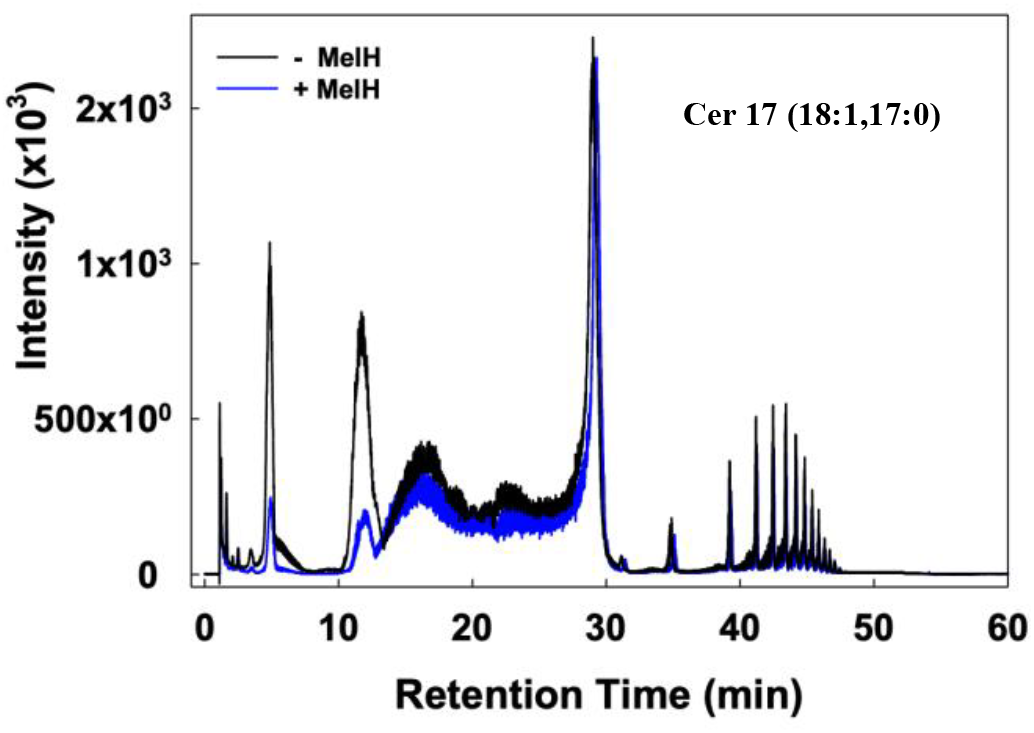
Relative abundance of the lipids from CHCl_3_:MeOH (85:15 v/v) fraction. The overlayed extracted ion chromatograms represent lipids from culture filtrates of the *melH* strain grown on propionate before and after addition of active MelH. A distinct reduction in ion intensity was observed after addition of MelH at retention times 5 mins and 10 mins. Ceramide17 (18:1, 17:0) is used as an internal standard.

To determine the molecular identities of the MelH-sensitive peaks, we used **SIRIUS** to predict best-fit molecular formulas and to evaluate the MS2 fragmentation patterns (14). Next, the data were processed with a custom Python script, sorting the m/z values with a mass accuracy of up to two decimal places (Fig 3). Only masses showing at least a two-fold change between pre- and post-treatment were considered for further analysis. The sorted masses were then analyzed with Bruker Metaboscape software to deconvolute adducts and to obtain accurate predicted molecular formulas. Most of the masses identified showed molecular formulas with 1–5 degrees of unsaturation and the presence of multiple oxygen atoms. The primary ions from both pre- and post-MelH treatment revealed several prominent peaks that include m/z values of 452.97, 604.47, 611.57, 629.10, 673.93, 714.47, 758.23 (Fig 4,5) (Table 1). The calculated molecular formulas ranged from C_60-100_H_160-200_O_3-4_, suggesting that these compounds belong to the class of Mycolic Acids (MA) (Table 1) (19). The observed fragmentation patterns closely resembled those of commercially available MA standards (Fig 6,7).

**Table 1:** Observed epoxide and diol masses with respective intensities in pre and post MelH treatment fractions with fold change.

| m/z | calculated<br>M | MF | Adducts | RT<br>(mins) | delta<br>m/z | unsaturation | fold<br>change | Diol<br>mH<sub>2</sub>/z-18 |
| --- | --- | --- | --- | --- | --- | --- | --- | --- |
| 452.97 | 902.89 | C61H122O3 | M+H+H | 3.21 | 0.07 | 1 | 6.8 | 452.27 |
| 604.47 | 1206.79 | C82H156O4 | M+H+H | 4.35 | 0.08 | 5 | 4.3 | 604.41 |
| 611.57 | 1221.15 | C83H158O4 | M+H+H | 3.25 | 0.15 | 5 | 5.8 | 611.63 |
| 629.11 | 1256.19 | C85H170O4 | M+H+H | 3.52 | 0.11 | 1 | 16.4 | 629.11 |
| 673.93 | 1345.89 | C92H176O4 | M+H+H | 4.55 | 0.04 | 5 | 12.3 | 673.93 |
| 714.47 | 1426.92 | C97H196O4 | M+H+H | 4.82 | 0.19 | 1 | 13.4 | 714.49 |
| 758.23 | 1515.15 | C104H200O4 | M+H+H | 5.11 | 0.11 | 5 | 12.4 | 758.23 |

\*\*The epoxide m/z represents the filtered masses that are changed upon addition of MelH with respective RT and fold change. The post treatment diol masses are represented as (mH<sub>2</sub>/z-18) since under LC conditions, the ions generated are usually a stable carbocation upon cleavage of one -OH group.

**Fig 3:**
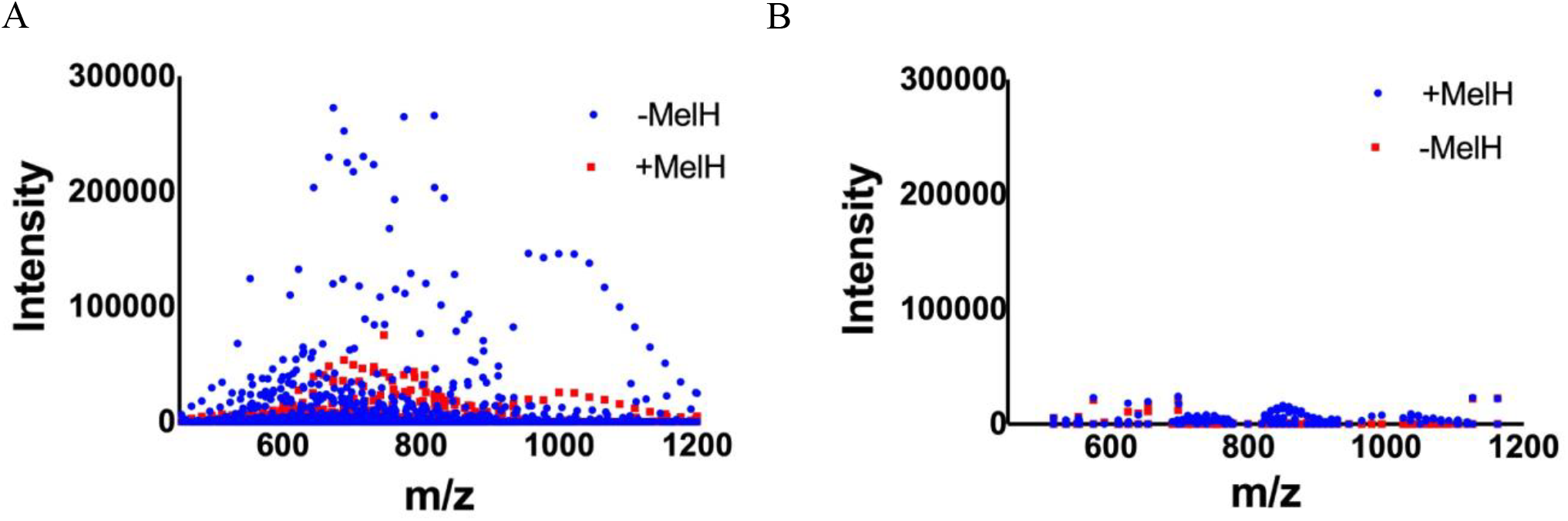
Ion Distribution Plots of m/z vs intensity. (A) Ion distribution plot vs intensity for all m/z observed between 3 and 12 mins The blue dots represent pre-MelH treatment ions whereas the red dots indicate post-MelH treatment ions. (B) Ion distribution plot vs intensity for all m/z observed between 20 and 25 mins. The pre/post-fold changes were determined and ions with a fold-change greater than 2 were selected for further analysis

**Fig 4:**
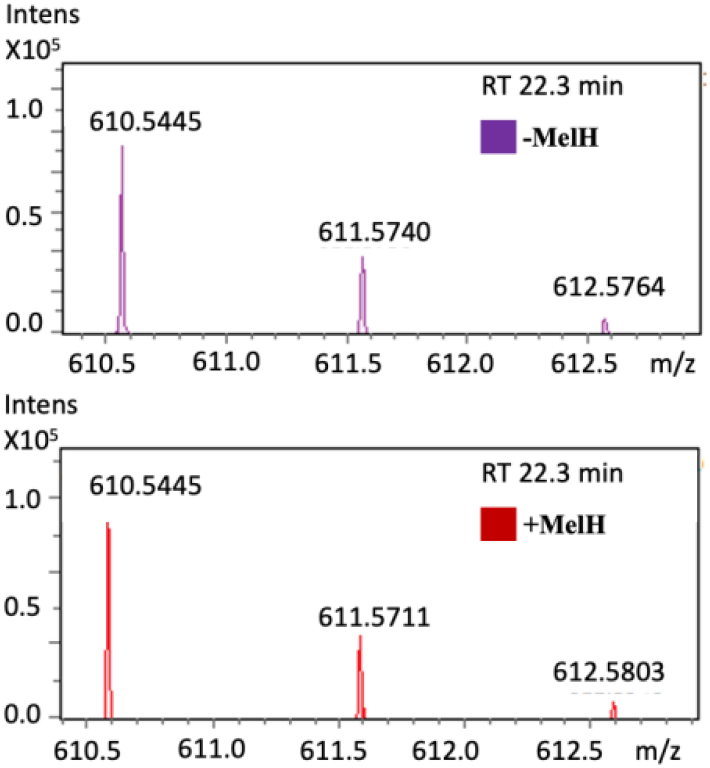
A. Positive control showing no apparent change in intensity for m/z 610.52 pre and post MelH treatment at RT 22.3 min. The ion is singly charged.

**Fig 5:**
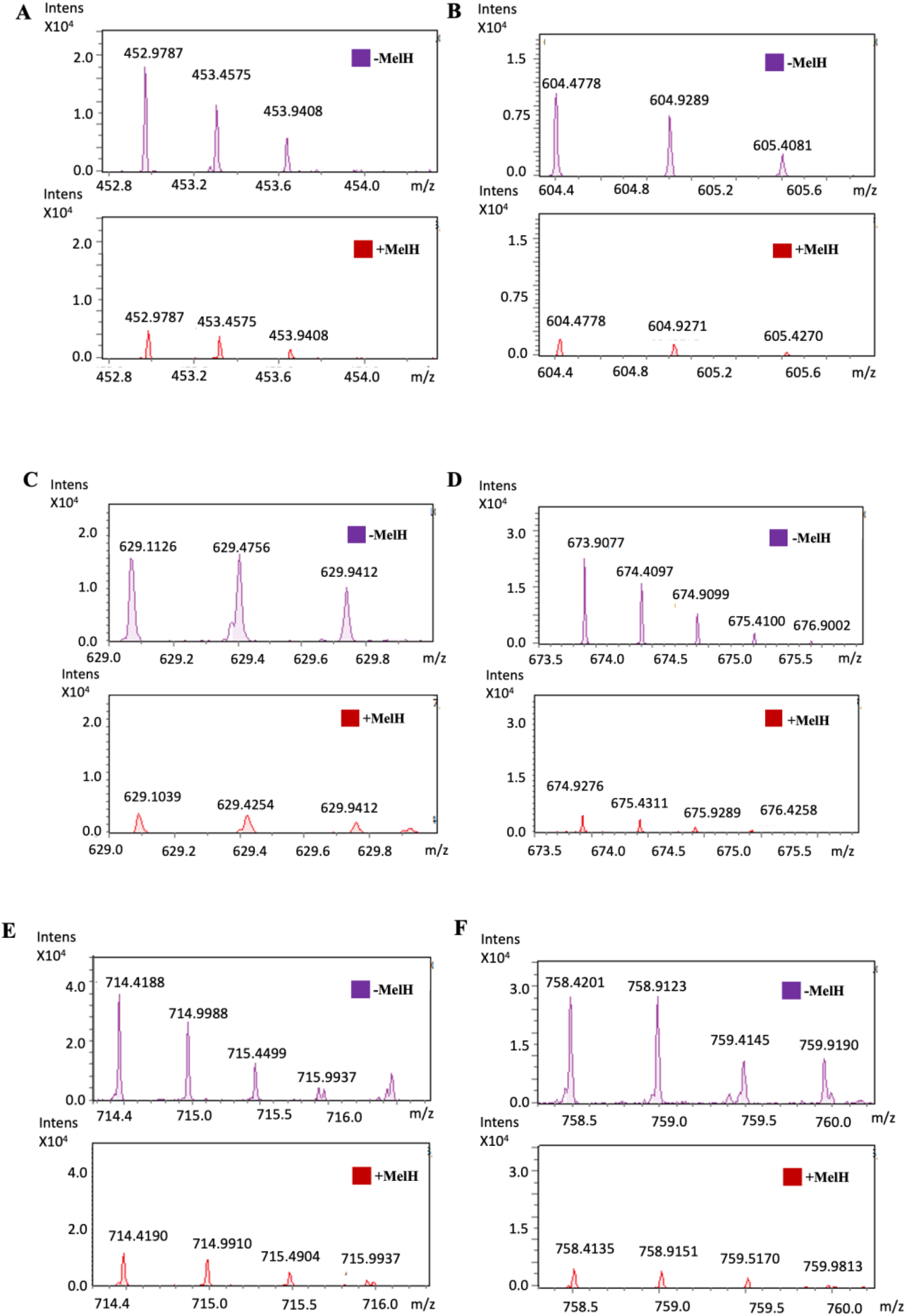
A, B, C, D, E, F represent the MS1 of individual m/z that are significantly changed after active MelH addition. The m/z’s are 452.97, 604.47, 629.10, 673.45, 714.41, 758.42. All ions are doubly charged.

**Fig 6:**
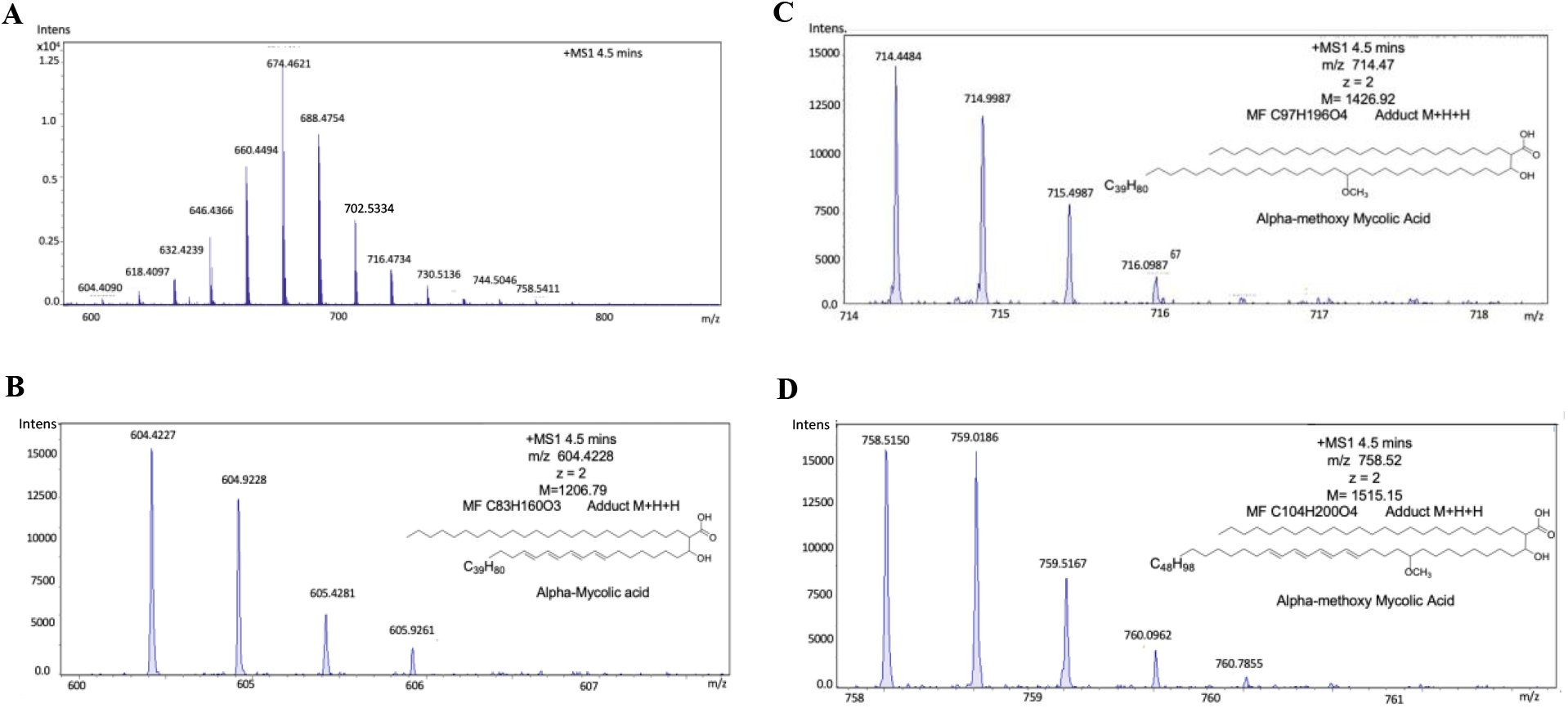
(A). MS1 spectrum of commercially available mycolic acid from *Mtb*. (B, C, D). m/z 604.4, 714.4 and 758.5 represent alpha and methoxy mycolic acids which eluted at RT 4.5 mins.

**Fig 7:**
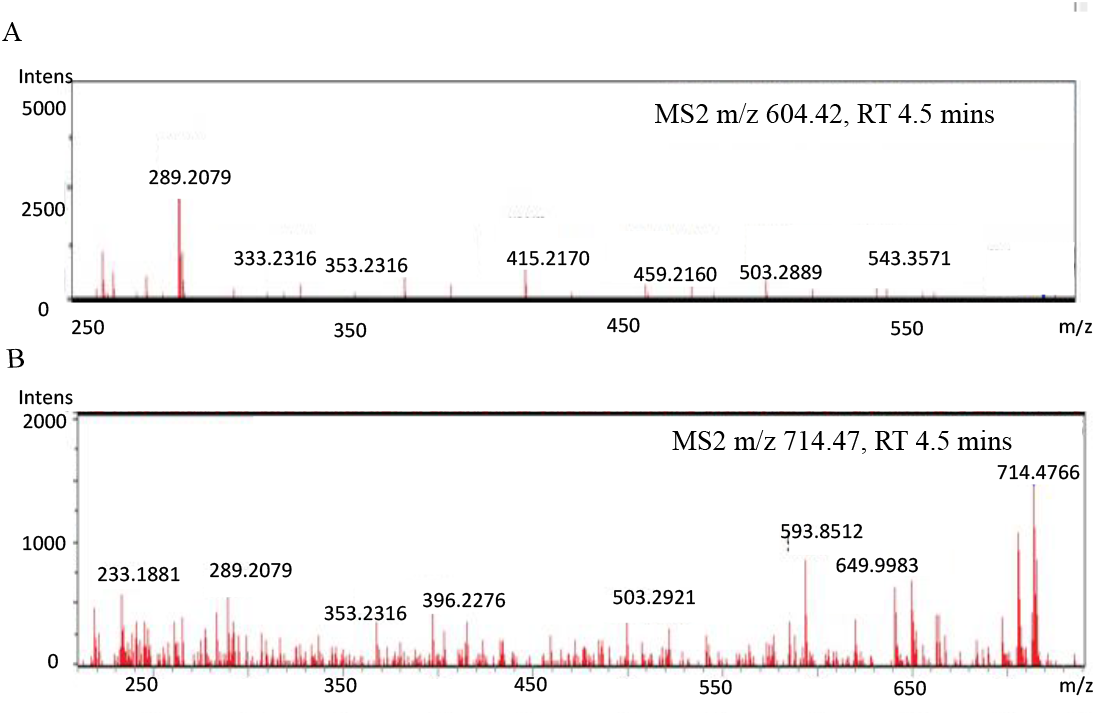
MS2 spectrum of commercially available mycolic acid standard extracted from *M. tuberculosis*. (A) MS2 of m/z 604.42, (b) MS2 of m/z 714.47.

Since diols may lose a hydroxyl group under LC-MS ionization conditions, the masses for the proposed parent diol mH_2_/z -18 were calculated. The masses include m/z 452.27, 604.41, 629.11, 611.63, 673.93, 714.49, 758.23. The corresponding diol masses obtained in the post MelH treatment fraction with individual intensities are presented in Table 2. The highest ion count was observed for m/z 714. 49, followed by m/z 758.23 which correspond to a parent diol mass of 1496.92 and 1515.45 respectively (Table 2). Although the position of the epoxide or the keto group cannot be determined from the data, the molecular formulas and the fragmentation pattern clearly suggest the presence of the epoxide or keto functional groups in the molecule.

**Table 2:**
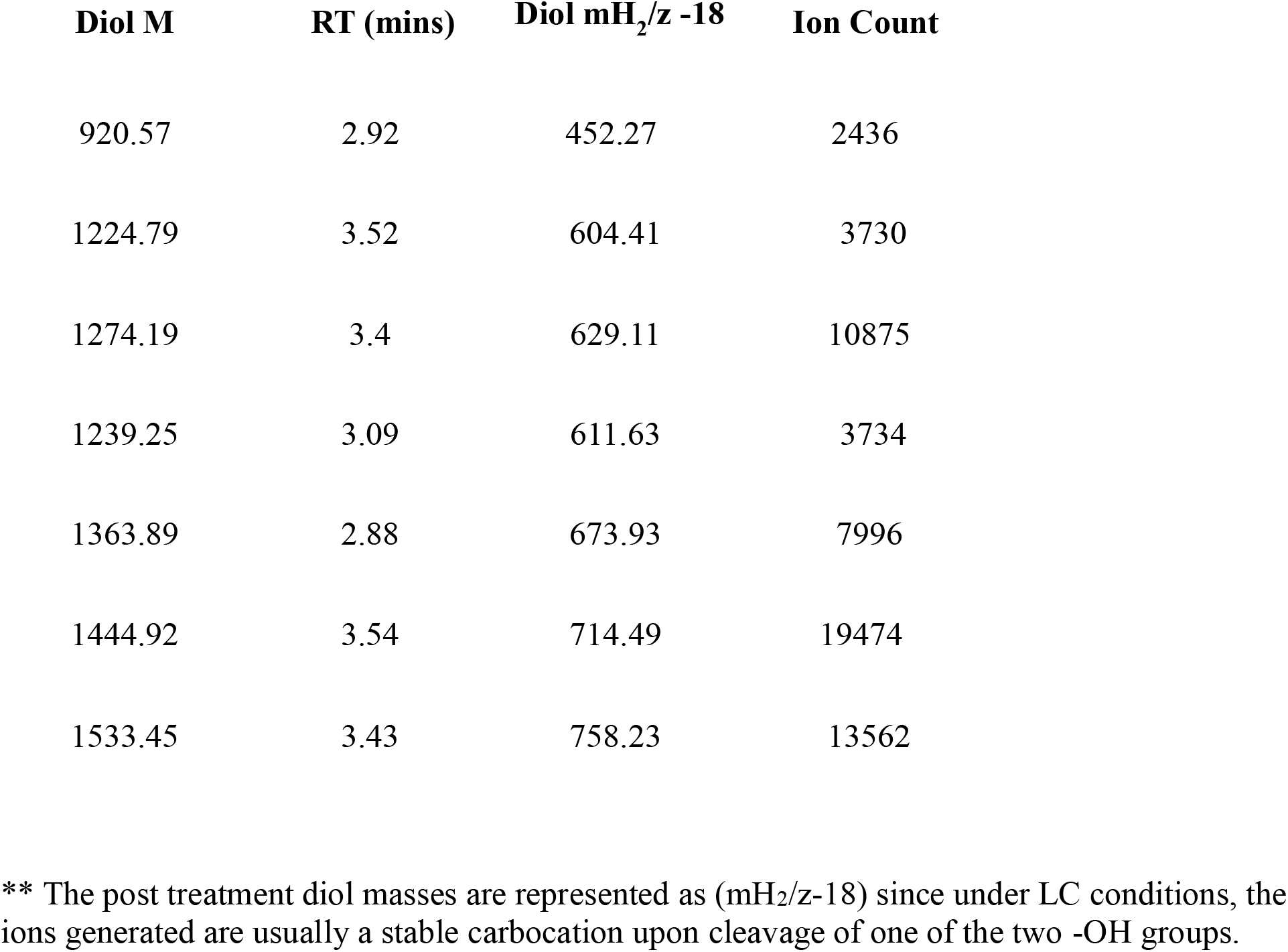
Respective diol (mH_2_-18) masses and Calculated Diol M with their intensities observed in post MelH trearment fraction:

| **Diol M** | **RT (mins)** | **Diol mH<sub>2</sub>/z -18** | **Ion Count** |
| --- | --- | --- | --- |
| 920.57 | 2.92 | 452.27 | 2436 |
| 1224.79 | 3.52 | 604.41 | 3730 |
| 1274.19 | 3.4 | 629.11 | 10875 |
| 1239.25 | 3.09 | 611.63 | 3734 |
| 1363.89 | 2.88 | 673.93 | 7996 |
| 1444.92 | 3.54 | 714.49 | 19474 |
| 1533.45 | 3.43 | 758.23 | 13562 |

\*\* The post treatment diol masses are represented as (mH<sub>2</sub>/z-18) since under LC conditions, the ions generated are usually a stable carbocation upon cleavage of one of the two -OH groups.

Using the m/z values and the parent masses, structures of the potential MA species were proposed (Fig. 8). The fragmentation patterns from the MS2 spectra were used to identify the cleavage of the bonds and the determination of the detailed structure of the MA class of lipids from the molecular formulas (Fig 9). The bonds that were cleaved in all the structures include the bond that breaks and forms meroaldehyde along with polar -COO bond . All MA’s contain a conserved chain attached to another chain that resembles all the modifications including unsaturation and functional groups (19). Therefore Fig 8 suggests the main fragments that were observed in the class of mycolic acid substrates for MelH, further confirming the structure. MA’s are found in several forms, including alpha mycolic acid, keto, epoxy, and methoxy mycolic acids (20). Since MelH is an epoxide hydrolase, we identified the corresponding diol species. Table 2 suggests the diols of the epoxide species identified and hence we can establish epoxide MA as the major substrate for MelH.

**Fig 8:**
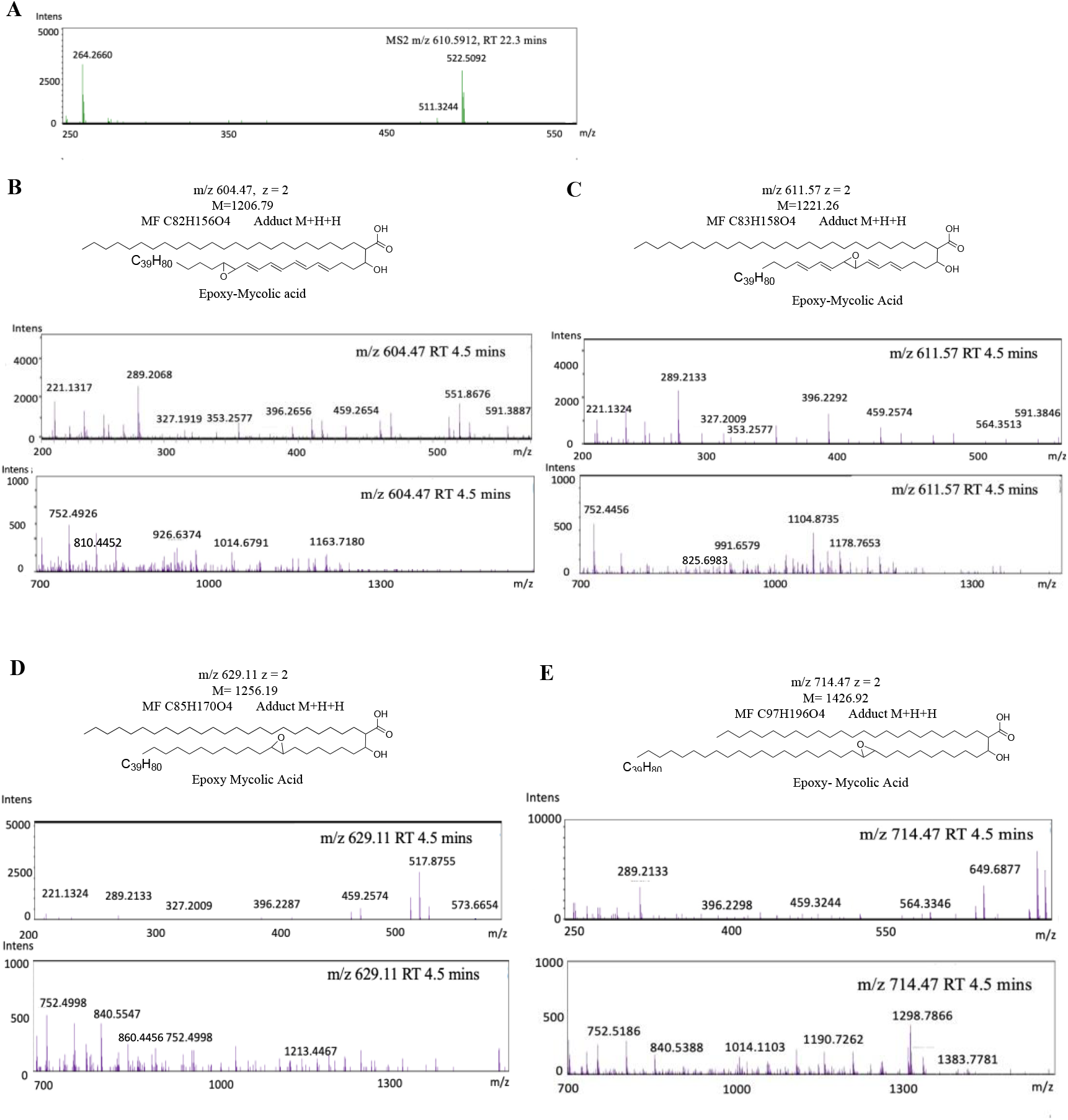
MS2 spectra for individual ions. (A). negative control MS2 spectrum of m/z 610.59. (B, C, D, E) MS2 spectra of the filtered ions with m/z 604.47, 611.57, 629.11 and 714.47 respectively. The mycolic acid structures are determined from the MS2 fragments of the individual ions.

**Fig 9:**
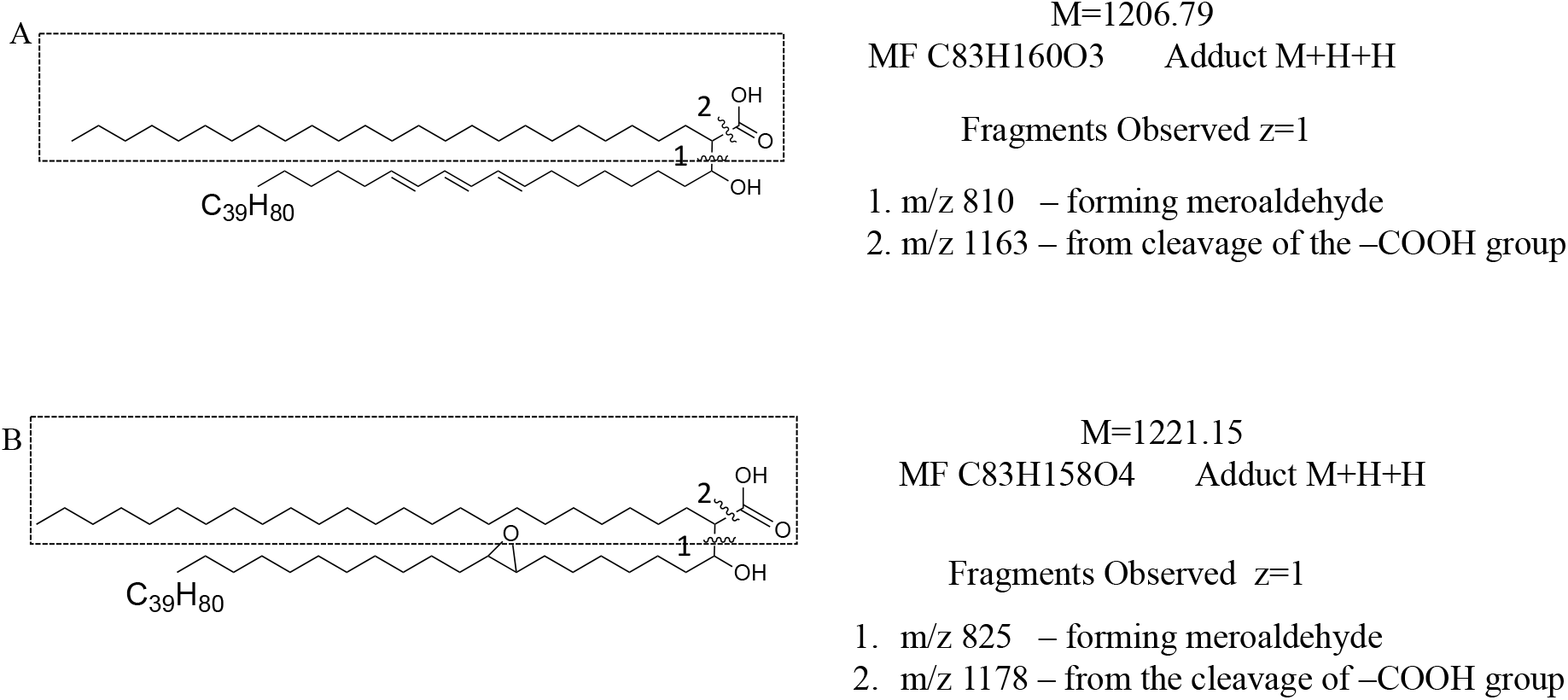
Proposed fragments corresponding to ion masses observed in MS2 spectra. The conserved cleavage pattern includes the bond forming meroaldehyde and carboxylic acids. (A) represents the representative cleavage points for alpha mycolic acids (B) represents the representative cleavage points for epoxy mycolic acids

## Discussion

So far, *melH* has been demonstrated to be involved in oxidative stress resistance during *Mtb* infection, but the physiological substrate of this protein remains unknown (9). In this study, we aim to identify the MelH substrate, which will also provide insights into MelH’s role in maintaining persistent infection and lipid homeostasis for sustained infection in *Mtb*.

To explore MelH’s role in catalyzing lipid metabolites, we first screened carbon sources where the deletion of MelH caused phenotypic defects (9). We identified glycerol, propionate, and cholesterol as key carbon sources for our experiments. We used *Mm WT* and *ΔmelH Mm* strains to demonstrate the effect of *melH* in vitro.

Initially, the competition assay suggested that when *Mm* is grown with propionate as a carbon source, lipids extracted from the culture filtrate of the *ΔmelH Mm* strain showed the highest abundance of a competitor (Fig. 1). Since only the culture filtrate lipid extracts showed accumulation of the physiological substrate, this suggests that the species might be toxic to the cell and could potentially be pumped out by transporters.

Therefore, total lipids extracted from the culture filtrate of the *ΔmelH Mm* strain were further fractionated using organic solvents (petroleum ether, chloroform [CHCl_3_], and an increasing gradient of methanol [MeOH]) to extract lipid classes, from the most hydrophobic to the least hydrophobic. For each fraction, active MelH was added, and both treated and untreated fractions were further subjected to LC-MS analysis. We observed a significant decrease in ion counts in the treated fraction of the CHCl_3_:MeOH (85:15 v/v) fraction compared to the untreated fraction.

Two distinct peaks were observed at RT 3-5 minutes and RT 10-12 minutes. Therefore, the masses from the RT 3-12 minute range were screened, and the best-fit molecular formulas were identified to predict the class of lipids that might act as a substrate for MelH. The molecular formulas suggested MA like lipids, and therefore the detailed MS2 fragmentation pattern was analyzed to predict the structure of this lipid class. MA’s are known to provide impermeability to antibiotics in the cell envelope of *Mtb*, thereby contributing to the natural resistance of mycobacteria against most antibiotics and playing a key role in mycobacterial virulence (21).

MA are a major, structurally complex long-chain fatty acids (C_60_–C_100_) that play a crucial role in the cell wall architecture of *Mycobacterium* species (22). They are thought to be synthesized in the cytoplasm and transported as trehalose esters via the MmpL3 transporter (23). MmpL3 is an established antibiotic target for *Mtb* (24). The biosynthesis of MA precursors involves two types of fatty acid synthases (FAS), FASI and FASII, which are both required to synthesize long-chain fatty acids known as meromycolic acids (25). *Mtb* on the other hand are known to synthesize several forms of MA’s namely, α-, methoxy-, keto-, and carboxy-mycolic acids with chain lengths varying from C_60_–C_100_ (26).

The chain length and abundance of MA depend on the growth rate and strain of the mycobacteria, with *M. smegmatis* having shorter chain lengths compared to *Mm* or *Mtb* (27). The virulence of mycobacteria is influenced by the structure of their MA’s, which include a meromycolic chain and an alpha branch (Fig. 6). Hence, MA’s can be divided into different classes based on the functional groups attached to the meromycolate. These classes are species-specific, and not all mycobacterial species contain all forms of MA’s (28). For example, keto and methoxy groups are found in pathogenic bacteria like *Mm* and *Mtb*, while the epoxy form is present in *M. smegmatis* but absent in *M. bovis* (29).

Since the post MelH treatment spectra showed a decrease in ion count than the pre MelH treatment spectra and upon considering that MelH is an epoxide hydrolase that catalyzes the conversion of an epoxide to a diol, we searched for epoxy mycolic acids in the *ΔmelH Mm* culture filtrate fraction. We found predicted epoxy mycolic acid structures for the sorted masses, which were confirmed with the MS2 fragmentation data (Table 1). This result suggests that MelH plays a role in detoxifying epoxides in pathogenic mycobacterial species like *Mm* and *Mtb*. The higher accumulation of epoxy mycolates in the *ΔmelH Mm* strain suggests that active MelH is responsible for converting epoxides to diols in the *WT*, as epoxy mycolates are not found in pathogenic mycobacteria (30).

Additionally, we also screened for potential diol products in our MS1 data (8,9,10). Although the best-fit molecular formulas for the masses were presumed to be epoxy or keto mycolic acids, the presence of diol products in the post-treatment spectra suggests that MelH is likely involved in converting epoxy-mycolic acids to their corresponding diols (Table 2). We also screened the masses in the *WT Mm* lipid extracts grown in propionate as a carbon source and they showed a higher ion count that in the *ΔmelH Mm*, which suggests that in the *WT*, diol products are of higher abundance than epoxy species because *Mtb* is a pathogenic mycobacterium and does not synthesize epoxy MA.

However, the commercially available mycolic acid standards represent only alpha and methoxy mycolic acids and did not represent epoxy species, as these mycolic acids were extracted through a method that involved alkaline hydrolysis, which cleaves epoxy and methoxy groups (31).

Overall, MA’s location in the inner leaflet of the bacterial outer membrane, forming a distinct layer called the mycomembrane and the essentiality of these lipid acids for viability and virulence makes them a potential drug target (32, 33, 34). Epoxy mycolic acids, in particular, play an important role as virulence factors, detoxification of which can increase the chances of sustained bacterial survival inside the host. Therefore, the role of MelH in catalyzing the conversion of epoxide MA to diols plays a crucial part in stress resistance during persistent and latent *Mtb* infection.

## Conclusion

In summary, we have identified a potential physiological substrate of MelH to be epoxidized mycolic acids through an untargeted lipidomic approach. These epoxidized mycolic acids are likely produced by ROS encountered by the mycobacteria. MelH converts the electrophilic epoxides to diol species which play an important role in detoxification of endogenous epoxides and can help promote persistent infection.

## ASSOCIATED CONTENT

### Supplementary materials

The following material is available online. Supplemental Tables, Figures and Spectra.

## AUTHOR INFORMATION

### Author Contributions

**Shreyoshi Chakraborti** — Program in Biochemistry and Structural Biology, Stony Brook University, Stony Brook, New York, 11794-5215 USA

**Jyothi S Sistla** — Department of Chemistry, Stony Brook University, Stony Brook, New York, 11794-3400

### Data availability

Lipidomics datasets from this study are deposited in the Global Natural Products Social Molecular Networking (GNPS; http://gnps.ucsd.edu). The data that support the findings of this study are available either within this article and its supplementary information files or upon reasonable request to the corresponding author.

## Acknowledgements

This work was supported by NIH grant R01AI134054. The lipidomic experiments were conducted at the Stony Brook CASDA Mass Spectrometry Center. The authors would like to thank Dr Nicole S Sampson for her invaluable input in review, conceptualization and funding acquisition. The authors would also like to thank Dr. Jeffrey D. Cirillo for generously providing the *ymm1 Mm WT* strain as a gift.

---

**Scheme 1:**
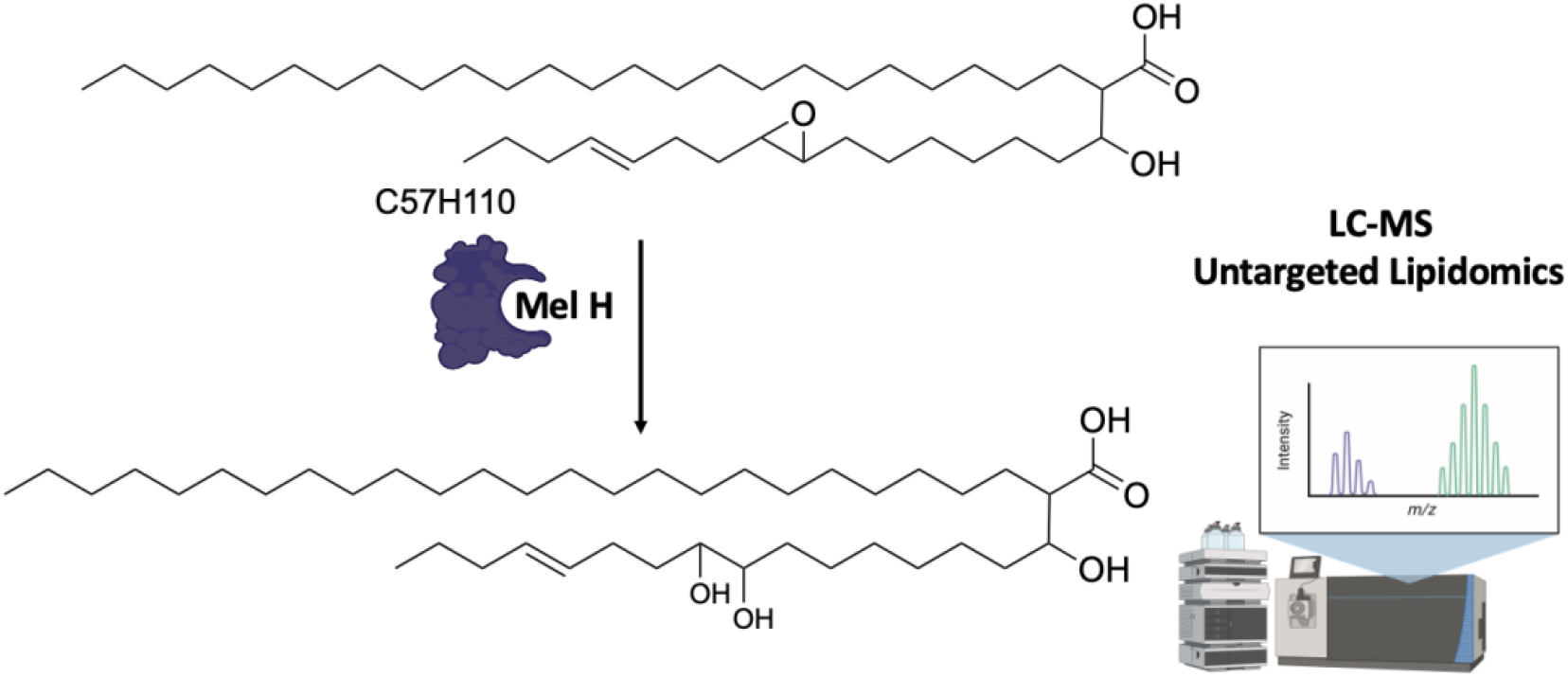
Schematic representation of MeIH reaction mechanism on epoxy MA (C27:0/C57:1) subtrates

