## Supplementary material for "Mycolic Acid like lipids act as substrates for *Mycobacterium tuberculosis melH*": Figure S1, Figure S2, Figure S3, Figure S4, Figure S5, Table S1, Table S2, Table S3

**Supporting Information for Publication:**

#### TABLE OF CONTENTS

**Figure S1:**. Plot of m/z vs fold change of the ions in RT 3-12 mins ................................2

**Figure S2:** Overlay plot from total lipid extraction of *Mm* WT and Δ*melH*..................... 3

**Figure S3:** Mass Spectra of commercially available Mycolic Acids ……………………..........4

**Figure S4:**. Total Ion Chromatograms from individual solvent reactions ……………….......5

**Figure S5:** MS1 spectra of masses after MelH treatment ……………………………………...6

**Table S1:** Substrate Epoxide masses observed in total lipid extracts from WT *Mm* …………..7

**Table S2:** Substrate Epoxide masses observed in total lipid extracts from KO *Mm* …………...8

**Table S3**: Product Diol masses observed in total lipid extracts from KO *Mm* …………..……...9

**Fig S1:**

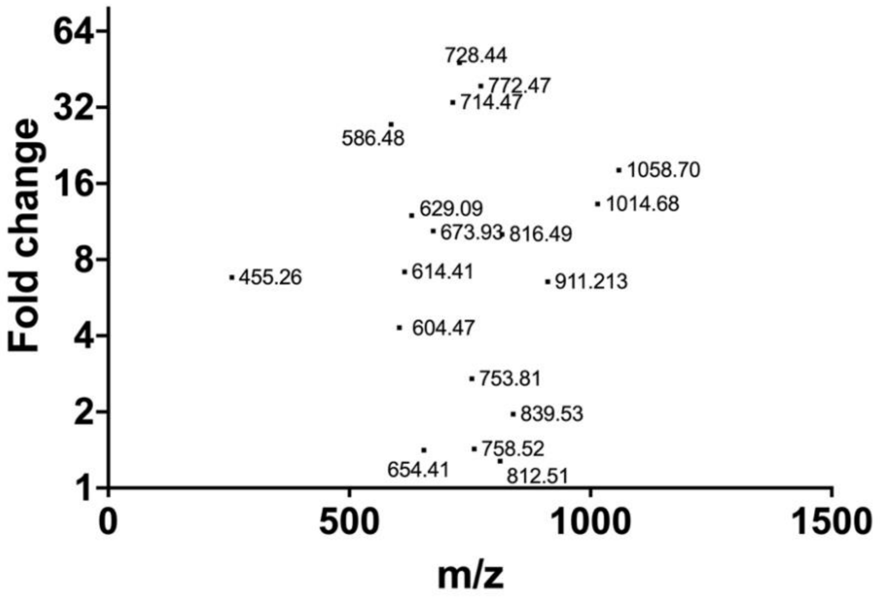


Figure S1: Plot of m/z vs fold change of the ions in RT 3-12 mins.

**Fig S2:**

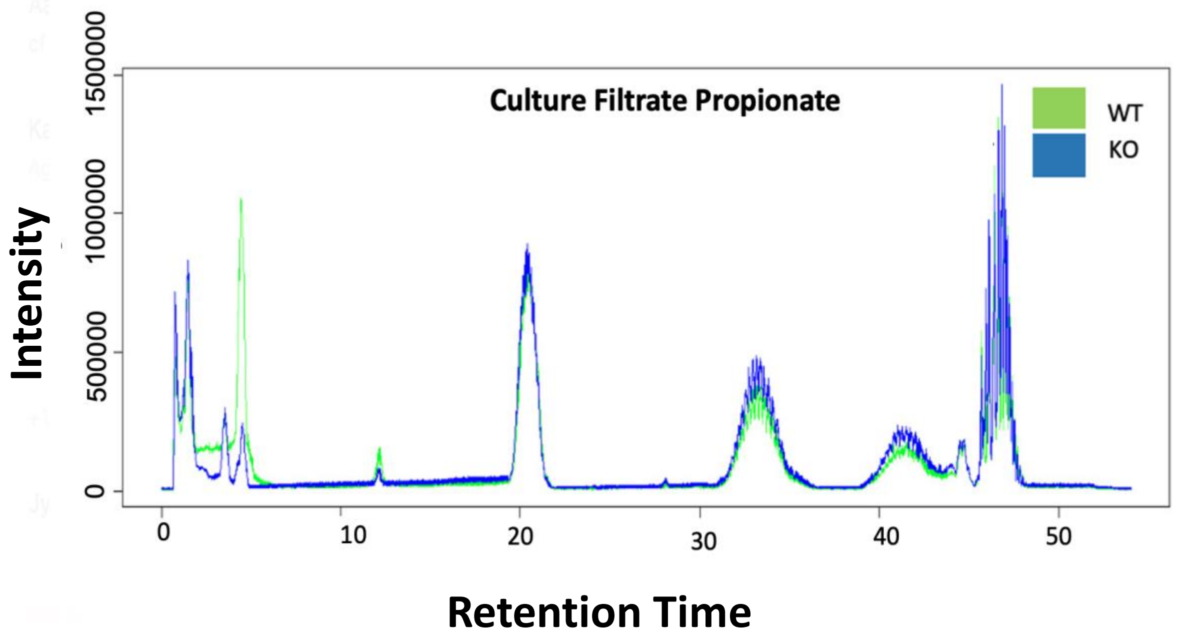


Figure S2: Overlay plot from total lipid extraction of culture filtrate grown in propionate from WT and Δ*melH*.

**Fig S3:**

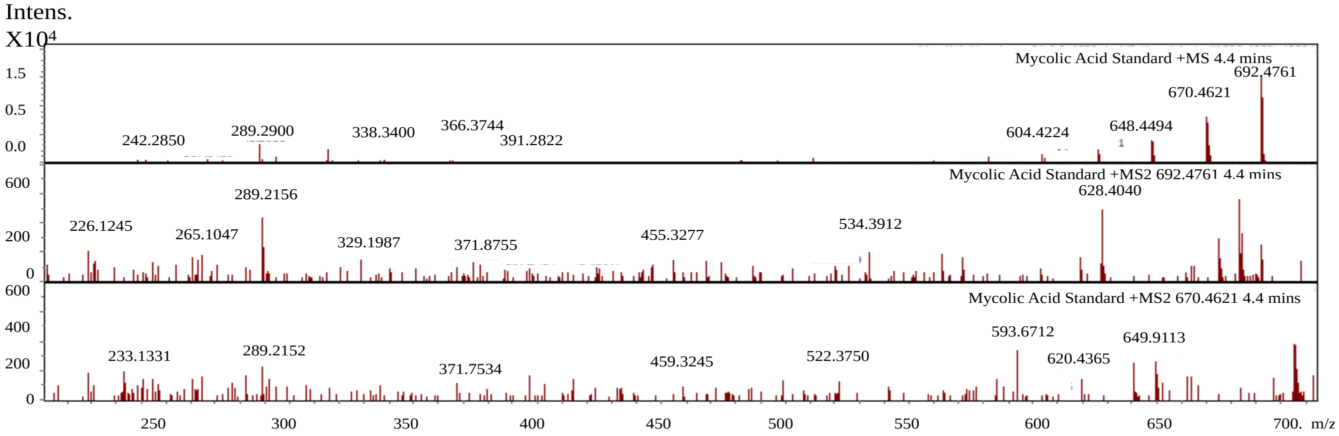


Fig S3: MS1 and MS2 spectra of commercially available Mycolic Acid Standard at RT 4.4 mins

### Fig S4:

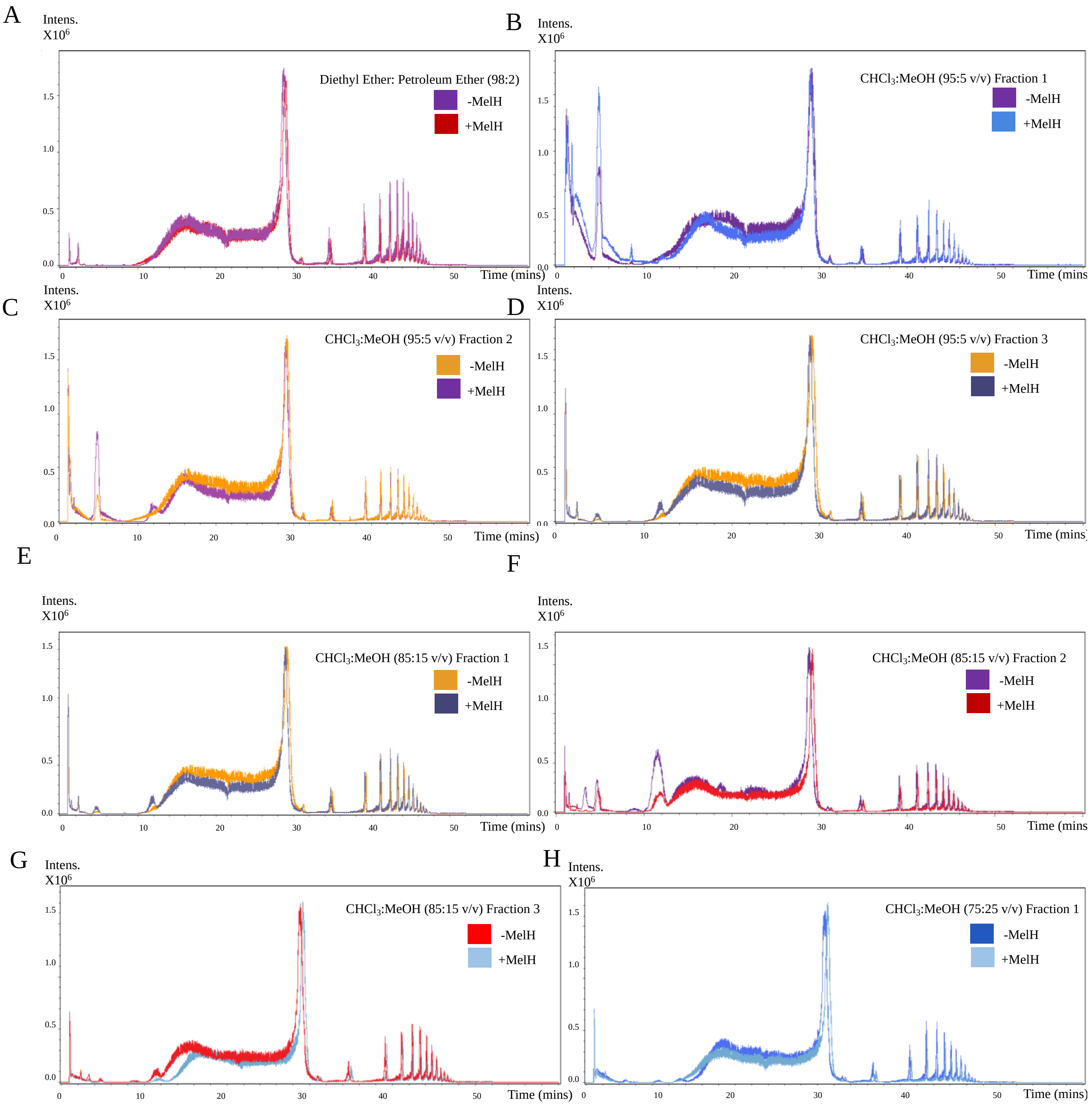

A
Intens. X10^6
Diethyl Ether: Petroleum Ether (98:2)
-MelH
+MelH
Time (mins)
B
Intens. X10^6
CHCl3:MeOH (95:5 v/v) Fraction 1
-MelH
+MelH
Time (mins
C
Intens. X10^6
CHCl3:MeOH (95:5 v/v) Fraction 2
-MelH
+MelH
Time (mins)
D
Intens. X10^6
CHCl3:MeOH (95:5 v/v) Fraction 3
-MelH
+MelH
Time (mins)
E
Intens. X10^6
CHCl3:MeOH (85:15 v/v) Fraction 1
-MelH
+MelH
Time (mins)
F
Intens. X10^6
CHCl3:MeOH (85:15 v/v) Fraction 2
-MelH
+MelH
Time (mins
G
Intens. X10^6
CHCl3:MeOH (85:15 v/v) Fraction 3
-MelH
+MelH
Time (mins)
H
Intens. X10^6
CHCl3:MeOH (75:25 v/v) Fraction 1
-MelH
+MelH
Time (mins)

Fig S4: Overlay total ion chromatograms of fractions from each solvent gradient. (A). Fraction from Diethyl Ether : Petroleum Ether (98:2) (B). CHCl3:MeOH (95:5) fraction 1 (C). CHCl3:MeOH (95:5) fraction 2 (D). CHCl3:MeOH (95:5) fraction 3 (E). CHCl3:MeOH (85:15) fraction 1(F). CHCl3:MeOH (85:15) fraction 2 (G). CHCl3:MeOH (85:15) fraction 3 (H). CHCl3:MeOH (75:25) fraction 1.

**Fig S5:**

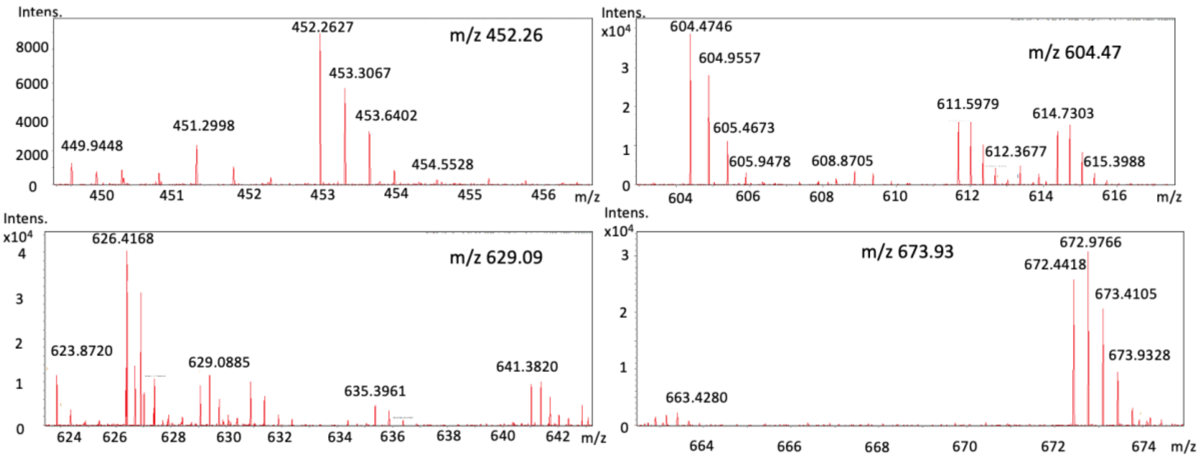


Fig S5: MS1 of m/z's that showed significant changes in intensity after active MelH addition. The m/z's shown here are 452.26, 604.47, 629.10, 672.98. All ions were doubly charged.

**Table S1: Observed WT MelH epoxide substrate masses found with respective intensities from Total Lipid Extraction from CF grown in propionate as a carbon source**

| Observed Mass WT | RT | m/z | Mean intensity of 3 replicates |
|---|---|---|---|
| 902.89 | 3.11 | 452.97 | 5267 |
| 1206.75 | 4.26 | 604.47 | 9877 |
| 1221.18 | 3.65 | 611.57 | 30098 |
| 1256.18 | 3.76 | 629.10 | 111280 |
| 1345.89 | 4.09 | 673.93 | 110098 |
| 1426.90 | 4.67 | 714.47 | 125298 |
| 1515.44 | 4.98 | 758.23 | 66345 |

**Table S2: Observed KO MelH epoxide substrate masses found with respective intensities from Total Lipid Extraction from CF grown in propionate as a carbon source**

| Observed Mass KO | RT | m/z | Mean intensity of 3 replicates |
|---|---|---|---|
| 902.89 | 3.11 | 452.97 | 6637 |
| 1206.74 | 4.26 | 604.47 | 8763 |
| 1221.19 | 3.65 | 611.57 | 5873 |
| 1256.19 | 3.76 | 629.10 | 13415 |
| 1345.89 | 4.09 | 673.93 | 1245 |
| 1426.91 | 4.67 | 714.47 | 6345 |
| 1515.44 | 4.98 | 758.23 | 5642 |

**Table S3: Observed KO MelH product diol masses found with respective intensities from Total Lipid Extraction from CF grown in propionate as a carbon source**

| Corresponding diol M | RT | Diol $mH_2/z$ -18 | Ion Count | Corresponding epoxy M |
|---|---|---|---|---|
| 1274.19 | 3.38 | 629.10 | 3562 | 1256.19 |
| 1239.25 | 3.11 | 611.63 | 3735 | 1221.19 |
| 1363.89 | 2.95 | 673.93 | 2454 | 1345.89 |
| 1444.92 | 3.32 | 714.47 | 3982 | 1426.91 |
| 1533.45 | 3.27 | 758.23 | 4533 | 1515.42 |

** The post treatment diol masses are represented as ($mH_2/z$-18) since under LC conditions, the ions generated are usually a stable carbocation upon cleavage of one of the two -OH groups.
